# Improving Variant Effect Prediction by Steering Sparse Mechanistic Features in Protein Language Models

**DOI:** 10.64898/2026.05.12.724472

**Authors:** Mingqing Wang, Meng Yuan, Athanasios V. Vasilakos, Yonghong He, Zhixiang Ren

## Abstract

Protein language models (PLMs) like the ESM series encapsulate immense evolutionary knowledge within their high-dimensional continuous embeddings. However, these latent representations are densely entangled, obscuring the fine-grained biophysical constraints necessary for precise functional resolution. To unlock the full expressive power of these embeddings, we propose PLM-SAE, a mechanistic framework that employs Sparse Autoencoders (SAEs) to disentangle PLM representations into discrete, biologically interpretable activations. By isolating and directly intervening on critical functional features, we fundamentally enhance the structural and mutational awareness of the underlying embeddings. We rigorously validate this embedding enhancement on variant effect prediction (VEP). In the unsupervised zero-shot setting, our sparse modulation elevates the state-of-the-art ESM-3 model, yielding performance improvements across 114 deep mutational scanning datasets and delivering an 80.8% relative improvement on challenging targets like the human E3 ubiquitin ligase HECD1. Furthermore, our target-specific differentiable gating mechanism achieves consistent performance gains in over 80% of evaluated datasets with an average Spearman ***ρ*** increase of +0.138. Finally, extending this approach to a cross-fitness multitask architecture establishes new state-of-the-art results on 17 VenusMutHub datasets, highlighted by a 169.0% performance surge in small-molecule binding predictions. Our work demonstrates that refining the highly entangled latent manifold via sparse modulation provides a robust and generalizable foundation for enhancing downstream PLM capabilities.

## 1 Introduction

Predicting the functional consequences of genetic variants, known as variant effect prediction (VEP), is a fundamental challenge in computational biology, with profound implications for understanding disease variants and evaluating protein fitness. Recently, protein language models (PLMs) have emerged as powerful tools for this task. By pre-training on large-scale evolutionary sequence databases, PLMs can capture complex mutational landscapes. Models such as ESM-1v and the recently introduced ESM-3 simulate evolutionary dynamics to enable zero-shot prediction of mutation effects [1, 2]. Other architectures, including ProSST and SaProt, incorporate structural awareness into language modeling to further refine these predictions [3, 4]. Furthermore, deep generative models and structure-based networks like EVE and AlphaMissense have significantly advanced our ability to perform proteome-wide missense variant effect prediction [5, 6].

To further enhance the predictive power of PLMs and improve variant effect prediction, recent research has explored various augmentation strategies. Some works have focused on integrating biophysical dynamics into the training process [7, 8]. Other approaches leverage text annotations or contrastive learning with biological features to enrich the learned representations across different modalities [9, 10]. Additionally, model compression and reverse distillation techniques have been proposed to consistently scale and compress the collective knowledge of massive PLMs into more efficient forms [11, 12]. These enhancements have collectively pushed the boundaries of VEP, allowing models to extract deeper biological insights from sequence data.

Despite these significant advancements, current zero-shot variant effect prediction paradigms suffer from a critical limitation: they typically collapse the complex, highdimensional representations of PLMs into a single scalar log-likelihood ratio (LLR) (**Figure 1a**). This dimensional collapse wastes the fine-grained biophysical features hidden within the high-dimensional latent manifold. Consequently, it often obscures the exact physical mechanisms—such as subtle steric clashes or the disruption of delicate hydrogen bond networks—that ultimately dictate a mutation’s functional consequence. To address the entangled “black box” nature of deep neural networks, sparse autoencoders (SAEs) have recently gained traction as a powerful tool for mechanistic interpretability [13, 14]. Pioneering studies have successfully applied SAEs to protein language models to uncover biologically interpretable features and map the mechanistic biology hidden within these complex representations [15–17].

**Fig. 1.**
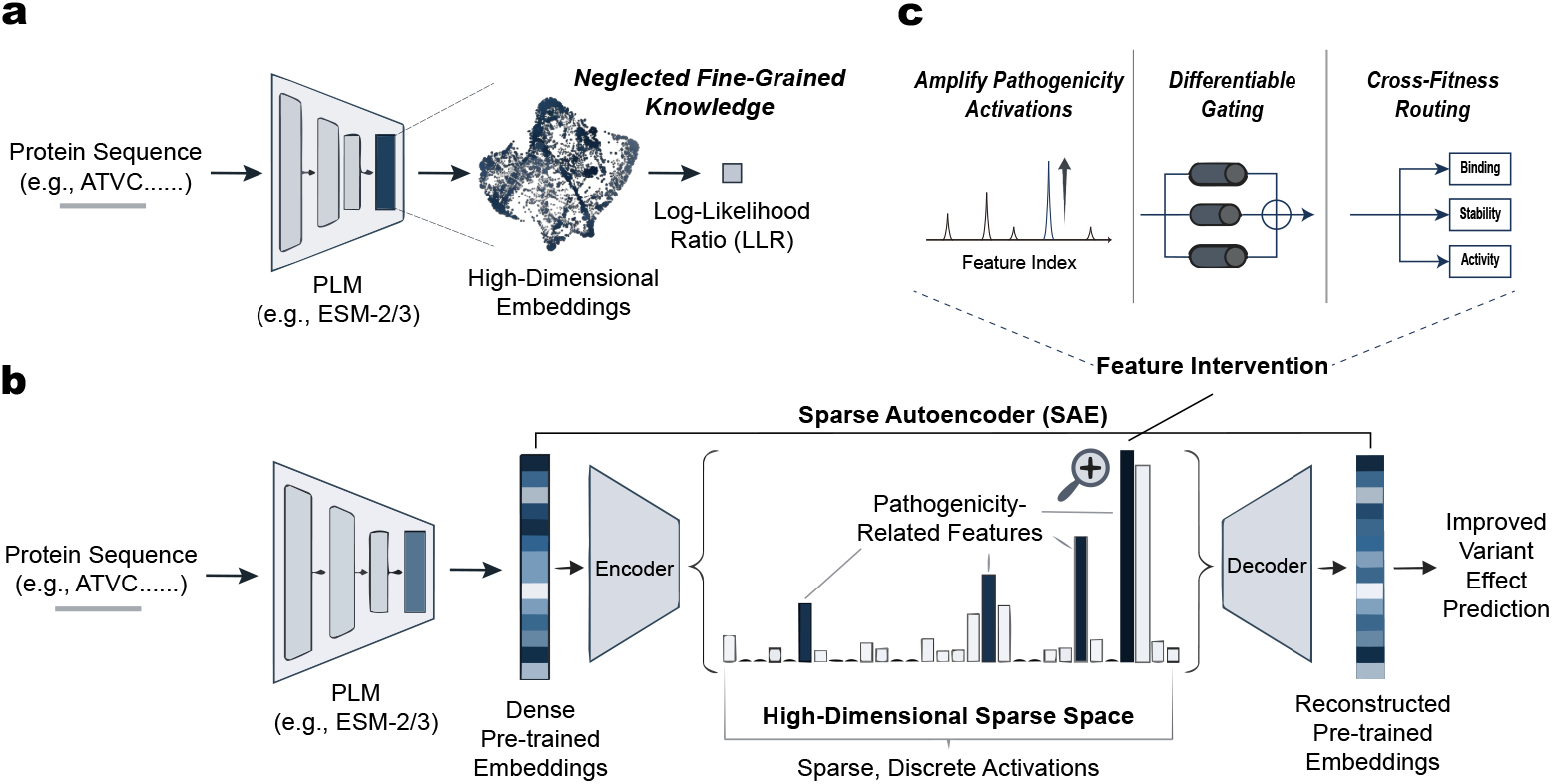
Representation decomposition and mechanistic intervention in protein language models. **a**, Limitations of the current zero-shot paradigm. Standard zero-shot variant effect prediction (VEP) typically collapses high-dimensional embeddings into a single scalar log-likelihood ratio (LLR). This bottleneck results in the neglect of fine-grained biophysical and evolutionary knowledge latent within the protein language model (PLM) manifold. **b**, SAE-based framework for knowledge extraction and intervention. We introduce a Sparse Autoencoder (SAE) to map dense pre-trained embeddings into a high-dimensional sparse space. This process yields sparse, discrete activations that disentangle complex representations into interpretable biological features. By identifying and magnifying pathogenicity-related features through direct intervention on these activations, the framework generates reconstructed embeddings that lead to improved variant effect prediction. **c**, Three distinct evaluation paradigms leveraging the disentangled functional activations. The framework systematically utilizes these features through unsupervised intervention by amplifying pathogenicity activations, target-specific supervised prediction via differentiable gating, and a multi-task architecture for cross-fitness routing across diverse biological properties such as binding, stability, and activity.

Building upon these interpretability efforts, we propose PLM-SAE, a novel representation enhancement framework that employs SAEs to disentangle continuous protein embeddings into discrete, biologically interpretable activations (**Figure 1b**). By explicitly identifying and modulating potentially pathogenic features within the sparse latent space, our approach effectively resolves dimensional collapse and fundamentally enhances the variant effect prediction capabilities of the underlying PLMs.

Crucially, our framework validates the enhanced quality of these modulated embeddings through three distinct paradigms (**Figure 1c**), demonstrating significant and measurable improvements:

- **Unsupervised Feature Amplification:** Even in the complete absence of target-specific priors, applying a simple element-wise boosting to SAE-extracted pathogenic features elevates the zero-shot capabilities of the state-of-the-art ESM-3 model. This unsupervised intervention improved predictions on 114 DMS datasets (52.5% win rate). Notably, it acts as a strong structural prior, providing a pronounced “rescue effect” on low-homology proteins with a 58.3% win rate and an average Spearman Δ*ρ* gain of +0.0129 against the baseline.
- **Target-Specific Differentiable Gating:** By introducing an end-to-end differentiable mask to dynamically route relevant sparse features, our supervised modulation generalizes exceptionally well, contrasting with the overfitting tendencies of standard linear probes (which degraded performance by an average Δ*ρ* of −0.033). Our targeted gating improved performance in over 80% of tested datasets (average Δ*ρ* = +0.138), achieving a 100% win rate in both the ‘Binding’ and ‘Virus’ categories with average improvements of +0.2346 and +0.1642, respectively.
- **Multi-Task Cross-Fitness Routing:** Shifting from isolated task modeling to a unified cross-fitness framework, our multi-task architecture leverages shared and private feature gates to establish new state-of-the-art benchmarks on 17 datasets within the VenusMutHub. This data diversity approach is particularly effective in complex domains, yielding a 169.0% relative improvement (from *ρ* = 0.1869 to 0.5028) in small-molecule binding tasks compared to the base ESM-2 model.

## 2 Results

### 2.1 Unsupervised extraction of pathogenic features enhances zero-shot variant effect prediction

Although protein language models (PLMs) learn complex evolutionary distributions during unsupervised pre-training, their knowledge is stored within dense, highly entangled embeddings. Current zero-shot prediction paradigms primarily rely on extracting the log-likelihood ratio (LLR) as a single scalar. This approach assumes that all biophysical constraints governing protein fitness can be seamlessly compressed into the marginal probabilities output by the final layer. However, this dimensional collapse often obscures the fine-grained physical mechanisms—such as subtle steric clashes or the disruption of delicate hydrogen bond networks—that dictate functional consequences. In the context of predicting variant pathogenicity, relying solely on these aggregated global probabilities limits our ability to resolve the exact structural perturbations causing a mutation’s deleterious effect.

To systematically uncover these hidden mechanisms, we repurpose the global likelihood not as a final prediction, but as a heuristic filter to isolate high-confidence pathogenic events. Specifically, we calculate the LLRs for all possible missense mutations within a dataset using a single base model (ESM-3). We then focus exclusively on the subset of variants exhibiting the lowest LLRs (Figure 2a, Steps 1 and 2). The rationale behind this selection is deeply rooted in the PLM’s pre-training objective: a severely depressed LLR indicates that the mutant amino acid fundamentally violates the evolutionary patterns and structural priors learned by the model. Consequently, these lower-LLR mutations serve as a highly enriched pool of potentially pathogenic or destabilizing variants.

**Fig. 2.**
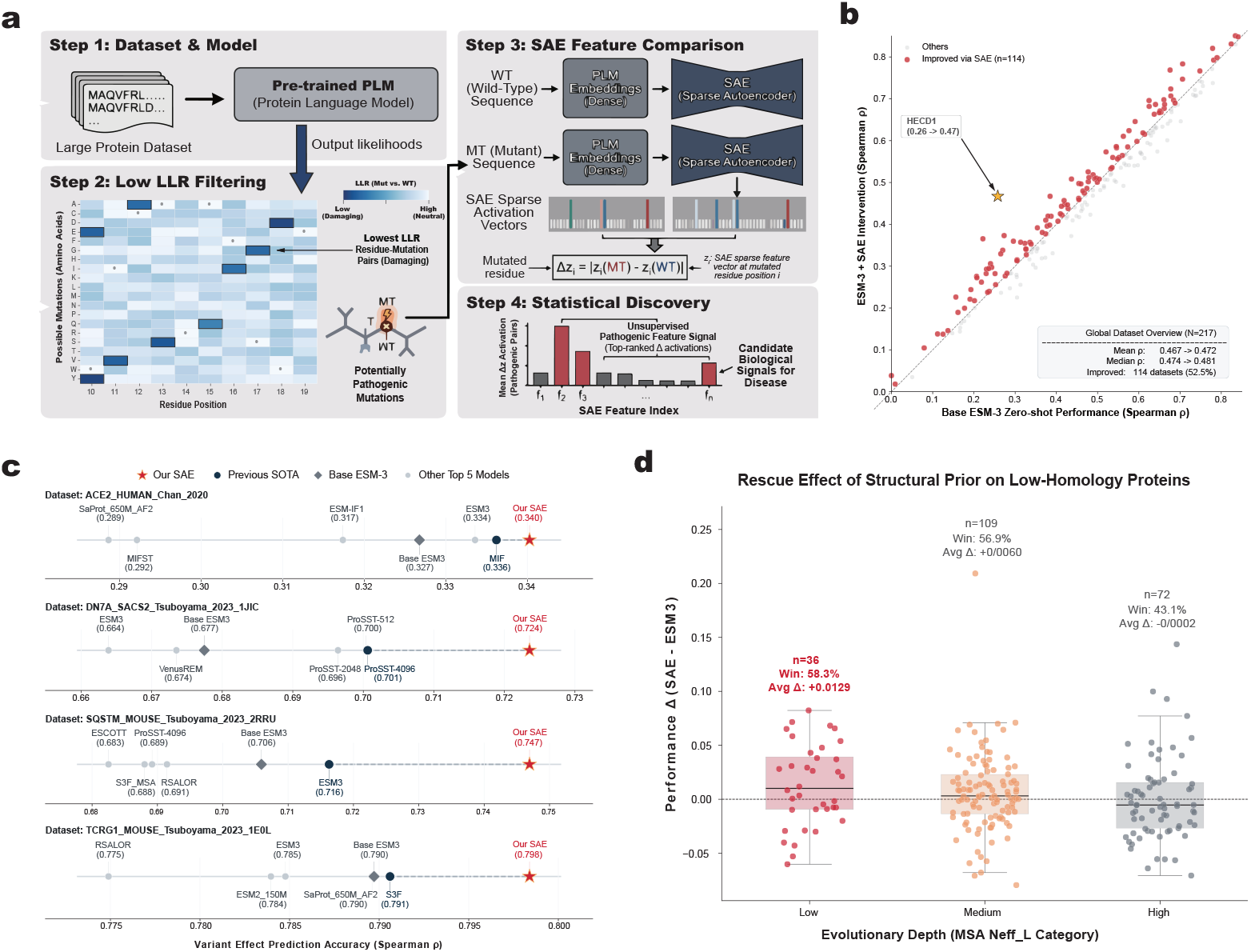
Unsupervised pathogenic feature discovery pipeline and evaluation of zero-shot variant effect prediction performance. **a**, The four-step pipeline for extracting and identifying pathogenic features using a Sparse Autoencoder (SAE). Steps 1 and 2 identify potentially pathogenic mutations by filtering for low log-likelihood ratios (LLRs) output by a pre-trained protein language model (PLM). Step 3 extracts dense embeddings for both wild-type (WT) and mutant (MT) sequences, mapping them to sparse activation vectors via the SAE to compute the activation difference (Δ*z*). Step 4 performs statistical aggregation of these differences across pathogenic pairs to discover candidate biological signals. **b**, Global performance comparison across deep mutational scanning (DMS) datasets (N=217). The scatter plot compares the zero-shot Spearman correlation (*ρ*) of the base ESM-3 model against the **ESM3-SAE-Zero** model. Red points denote datasets where the unsupervised intervention improved performance (114 datasets, 52.5%). **c**, Head-to-head performance evaluation on four representative DMS datasets. The **ESM3-SAE-Zero** model (red star) consistently outperforms the base ESM-3 (grey diamond), the previous state-of-the-art (SOTA, dark blue dot), and other top-tier models. **d**, Performance improvement (Δ Spearman *ρ*) stratified by evolutionary depth (MSA Neff_L categories). The **ESM3-SAE-Zero** intervention demonstrates a pronounced “rescue effect” on low-homology proteins, yielding the highest win rate (58.3%) and average performance gain (+0.0129), suggesting that the extracted features successfully compensate for the lack of evolutionary information by providing structural or biophysical priors.

Having captured these potential pathogenic mutations, we use them as specific “probes” to interrogate the model’s internal manifold. We introduce the **ESM3-SAE-Zero** model, which utilizes a sparse autoencoder (SAE) to project the continuous hidden representations of both the wild-type (WT) and the mutant (MT) sequences for these specific variants into a high-dimensional sparse space. In this disentangled space, biophysical properties are isolated into discrete activating neurons. By computing the activation difference Δ*z*_*i*_ = |*z*_*i*_(MT) − *z*_*i*_(WT)| across this filtered subset of pathogenic pairs, we can precisely pinpoint the specific feature dimensions that are most acutely perturbed (Figure 2a, Steps 3 and 4). Unlike relying on the ambiguous LLR scalar, **ESM3-SAE-Zero** utilizes the extremes of the LLR distribution to anchor our search, enabling the unsupervised statistical discovery of the exact mechanistic features—such as spatial conflicts or hydrophobic core disruptions—responsible for the pathogenicity.

To validate the efficacy of these unsupervised extracted features, we implemented a direct feature intervention strategy: after identifying the feature dimensions with the largest Δ*z*, **ESM3-SAE-Zero** amplifies their activation values to twice their original magnitude. We conducted systematic zero-shot evaluations across 217 deep mutational scanning (DMS) datasets from ProteinGym [18] (Figure 2b). The results demonstrated that this straightforward intervention improved performance on 114 datasets (yielding a win rate of 52.5%) and increased the overall average Spearman correlation (*ρ*) from 0.467 to 0.472. Most notably, this approach exhibited a profound rescue effect, successfully rectifying the exceptionally poor zero-shot predictions of the base ESM-3 model on specific proteins. For instance, on the human E3 ubiquitin ligase HECD1 dataset, the base ESM-3 model achieved a sub-par Spearman *ρ* of 0.26. Following intervention via **ESM3-SAE-Zero**, the performance surged to 0.47, representing an 80.8% relative improvement. This suggests that by amplifying the specific features isolated through our low-LLR probing strategy, the model can recover critical pathological signals that were previously buried in the dense embeddings.

Surprisingly, even though this framework relies on a singular, unsupervised feature enhancement step—without any task-specific supervision or complex postprocessing—it established new state-of-the-art benchmarks on several highly competitive datasets. On four representative DMS datasets (ACE2_HUMAN, DN7A_SACS2, SQSTM_MOUSE, and TCRG1_MOUSE), **ESM3-SAE-Zero** comprehensively outperformed all existing top-tier models (Figure 2c). This includes models explicitly optimized for structure (such as MIF-ST, which scored 0.292 versus our 0.340 on ACE2), models leveraging massive multiple sequence alignments (like S3F), and the latest advanced pre-trained models like ProSST. By isolating and intervening on mechanistic features, **ESM3-SAE-Zero** consistently surpassed both the base ESM-3 and the leading zero-shot competitors.

To further investigate the biological underpinnings of these performance gains, we stratified the results by evolutionary depth using the MSA Neff_L categories (Figure 2d). This analysis revealed a compelling pattern: the **ESM3-SAE-Zero** intervention yielded the most substantial improvements on low-homology proteins (achieving a 58.3% win rate and an average Δ*ρ* of +0.0129), with the relative gains gradually converging as evolutionary depth increased. From a biophysical perspective, PLMs struggle to infer fitness landscapes for proteins with low Neff_L values due to the scarcity of co-evolutionary signals. In these scenarios, the SAE-extracted features act as universal biophysical priors, encoding fundamental rules such as steric hindrance and hydrophobic core stability. This rescue effect indicates that our framework does not merely overfit to evolutionary noise; rather, it disentangles and learns generalizable protein structure-function relationships based on physical principles. By effectively compensating for the absence of evolutionary scale information, this intervention endows the language model with robust predictive capabilities even in evolutionary “dark zones.”

### 2.2 Differentiable feature masking identifies target-specific biological signals

Beyond the unsupervised statistical discovery of pathogenic features, we sought to determine whether the identification of critical biological signals could be further refined through an automated, end-to-end learning process. While unsupervised methods are effective at capturing general structural perturbations, different biological targets—each with unique structural environments and functional constraints—may rely on distinct subsets of the latent manifold. To explore this, we introduced the **ESM3-SAE-Target** model, which utilizes a differentiable feature masking mechanism designed to automatically isolate target-specific features from the sparse latent space of the protein language model (PLM) (Figure 3a).

**Fig. 3.**
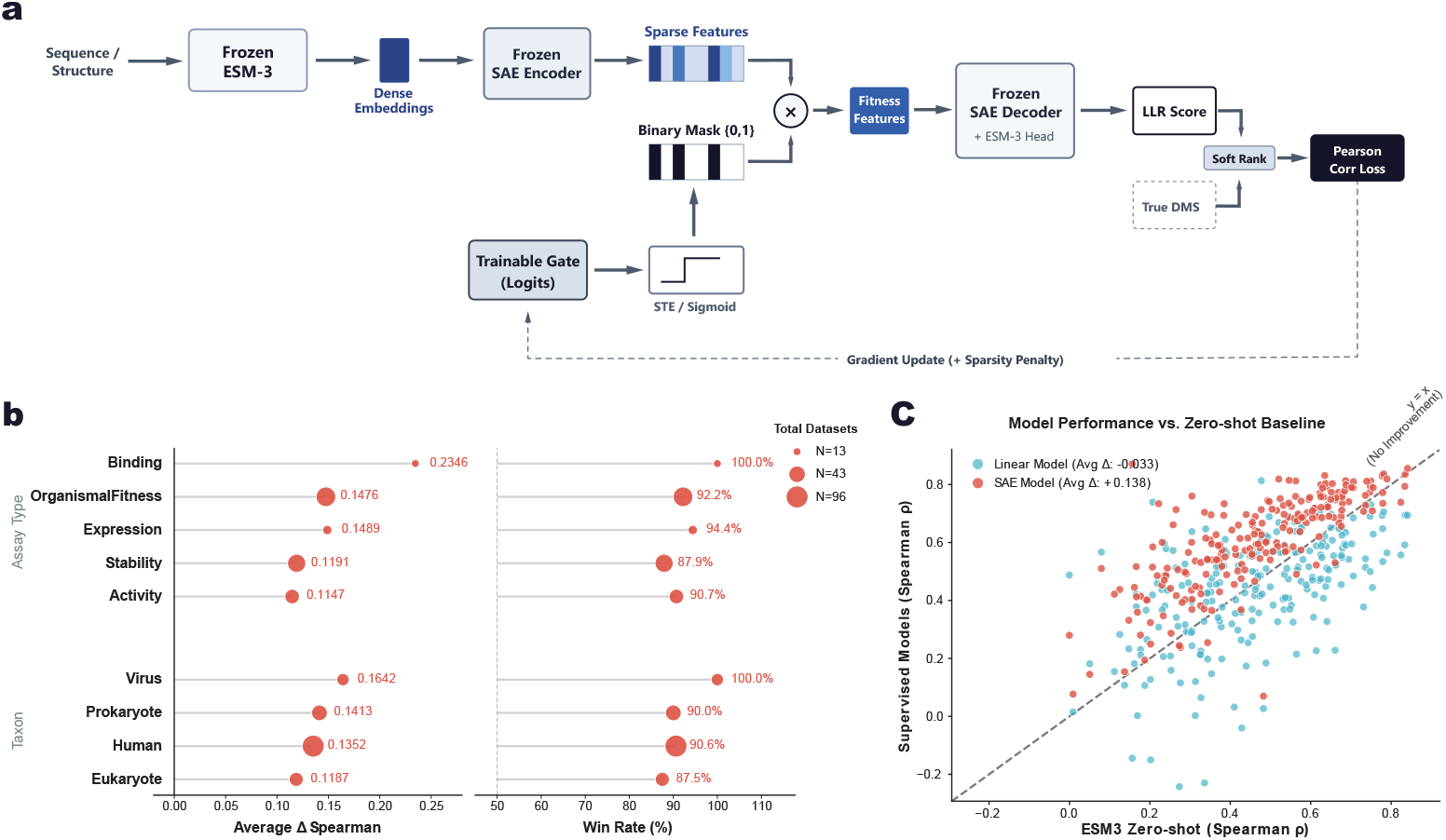
Target-specific feature selection via differentiable masking and supervised performance evaluation. **a**, Architecture of the supervised feature gating mechanism. Protein sequences are processed by a frozen ESM-3 and mapped to a sparse latent space via a frozen SAE encoder. A trainable gate, optimized using a Straight-Through Estimator (STE) with a sparsity penalty, generates a binary mask ({0, 1}) to isolate target-relevant fitness features. These selected features are reconstructed by the SAE decoder and projected by the ESM-3 head to compute log-likelihood ratio (LLR) scores. The gating module is trained using a soft-rank Pearson correlation loss against true deep mutational scanning (DMS) values. **b**, Performance breakdown of the **ESM3-SAE-Target** model across diverse assay types (e.g., Binding, Expression) and taxonomic categories. The charts report the average performance gain (Δ Spearman *ρ*) and win rate (%) relative to the zero-shot baseline, highlighting robust improvements across biological domains. **c**, Global comparison of supervised models versus the zero-shot baseline. The scatter plot demonstrates that **ESM3-SAE-Target** (Avg Δ: +0.138) effectively leverages supervision to consistently improve prediction accuracy across 96 datasets, whereas a standard linear probe (Avg Δ: −0.033) degrades performance, likely due to overfitting on the entangled continuous representations.

We evaluated this supervised gating framework using the comprehensive Deep Mutational Scanning (DMS) datasets from ProteinGym, adhering to the official five-fold random split protocol to ensure rigorous cross-validation and prevent data leakage. The core of **ESM3-SAE-Target** involves a trainable gating module that operates on the frozen SAE latent space. By employing a Straight-Through Estimator (STE) with a sparsity penalty, the model learns to generate a binary mask ({0, 1}) that selects only the most informative fitness-related features for a given protein assay (Figure 3a). This allows the framework to transition from a general mechanistic search to a targeted intervention tailored to the unique structural and functional requirements of the specific protein target.

The deployment of **ESM3-SAE-Target** resulted in widespread and substantial performance gains across a broad range of taxonomic groups and fitness types (Figure 3b). We observed consistent improvements in over 80% of the total datasets, with an average Spearman *ρ* increase exceeding 0.1 relative to the zero-shot baseline. Notably, the framework achieved a 100% win rate for all datasets within the ‘Binding’ fitness category and the ‘Virus’ taxonomic group, yielding remarkable average improvements of +0.2346 and +0.1642, respectively (Figure 3b). Such pronounced gains in binding assays suggest that the SAE effectively disentangles the specific geometric and electrostatic features essential for molecular recognition, which the supervised gate can then accurately identify and leverage for the specific target. Even in categories with lower average gains, such as ‘Stability’ (+0.1191) and ‘Eukaryote’ (+0.1187), the win rates remained exceptionally high (87.9% and 87.5%, respectively), demonstrating the robustness of the feature-masking approach.

To further validate the superiority of our method, we contrasted its performance with the conventional linear probing approach, which is frequently used to evaluate the quality of pre-trained representations. We found that standard linear probes, which fit a linear layer directly on dense embeddings, are highly prone to overfitting the training distribution, often resulting in a significant degradation of performance on unseen test sets (average Δ*ρ* = −0.033; Figure 3c). In contrast, **ESM3-SAE-Target** (average Δ*ρ* = +0.138) demonstrated superior generalization capabilities. By restricting the model to a sparse selection of physically-grounded latent features rather than allowing it to unrestrictedly weight the entire dense representation, our framework maintains the structural integrity of the pre-trained knowledge while successfully adapting to target-specific biological signals. This suggests that the “true” functional landscape is more accurately captured by a sparse selection of mechanistic features than by a complex linear combination of entangled embeddings, confirming that sparsity serves as a crucial regularizer for capturing authentic biological fitness.

### 2.3 Multi-task architecture for cross-fitness variant effect prediction

To further extend the utility of mechanistic intervention, we propose **ESM2-SAE-Cross**, a novel paradigm shifting from isolated task modeling to a unified cross-fitness framework. In this architecture, a single model is trained simultaneously on a diverse pool of fitness data, enabling joint feature discovery across all categories and performing inference through a modularized gating system (Figure 4a). To rigorously evaluate this multi-task capability, we utilized the VenusMutHub benchmark [19], which provides a comprehensive categorization of protein fitness into five distinct classes: *activity, ppi_binding, selectivity, small_molecule_binding*, and *stability*. Given that VenusMutHub primarily comprises sequence-only datasets without high-resolution structural information, we explicitly employed the ESM-2 model (650M) as our backbone for these experiments, demonstrating the adaptability of the SAE framework across different protein language model (PLM) architectures.

**Fig. 4.**
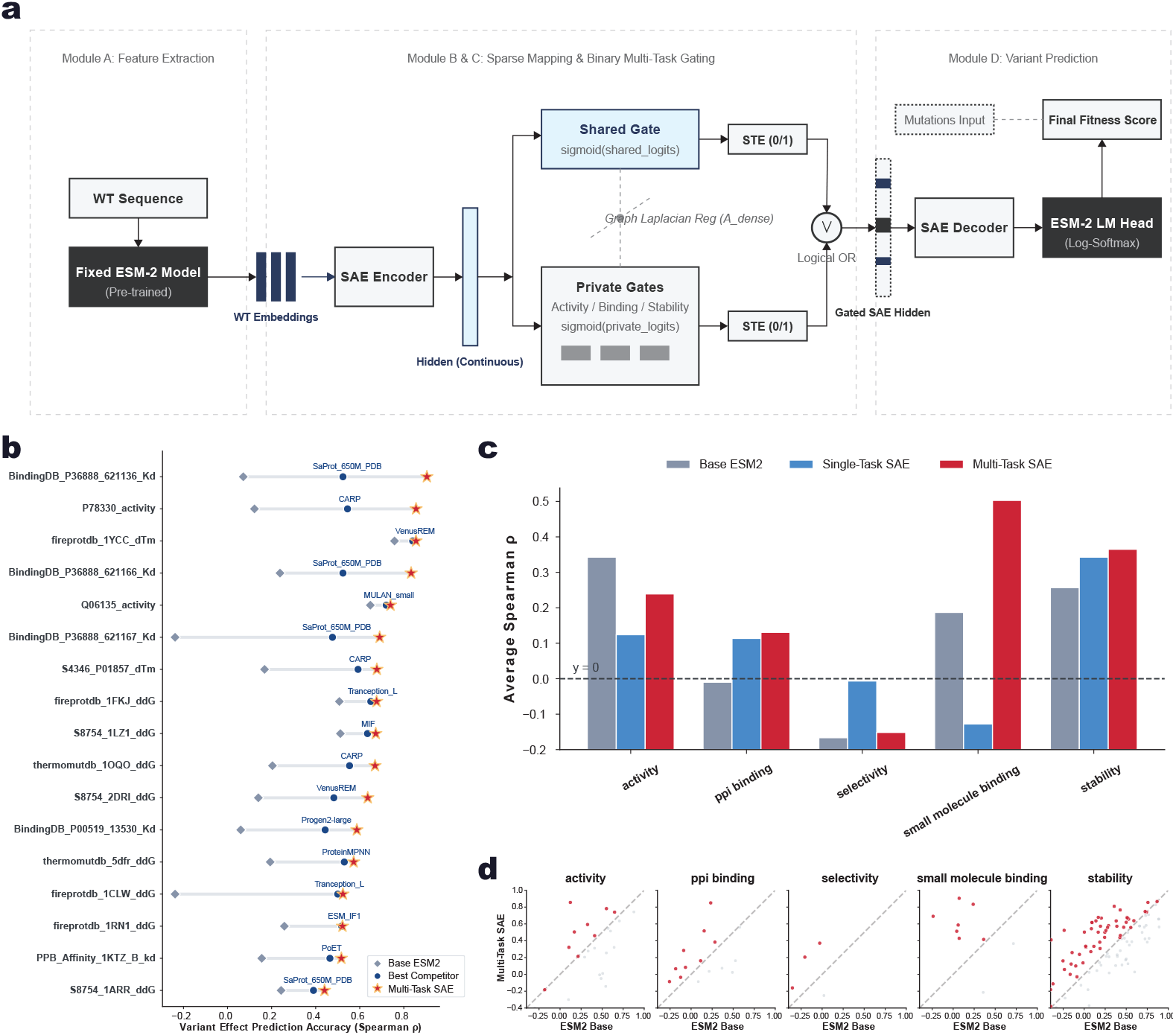
Multi-task feature gating architecture and performance evaluation on diverse variant effect prediction tasks. **a**, Structure of the multi-task SAE framework. Module A extracts embeddings using a fixed pre-trained ESM-2 model. Modules B and C perform sparse mapping and binary multi-task gating, utilizing an SAE encoder. The gating mechanism employs a shared gate and task-specific private gates (e.g., Activity, Binding, Stability) optimized via Straight-Through Estimator (STE). A Graph Laplacian regularization (*A*_*dense*) is applied to the shared logits. The gated hidden representations are combined via a logical OR operation and passed to the SAE decoder. In Module D, the ESM-2 language model (LM) head processes the mutations input alongside the reconstructed features to compute the final fitness score. **b**, Detailed performance comparison on individual deep mutational scanning datasets. **ESM2-SAE-Cross** (red stars) is evaluated against the Base ESM2 (grey diamonds) and the Best Competitor (blue dots) for specific datasets, measured by Variant Effect Prediction Accuracy (Spearman *ρ*). **c**, Bar plot summarizing the average Spearman *ρ* across five functional categories: activity, ppi_binding, selectivity, small_molecule_binding, and stability. **ESM2-SAE-Cross** consistently demonstrates improved performance compared to both the Base ESM2 and the single-task control model (**ESM2-SAE-Task**). **d**, Dataset-level scatter plots comparing **ESM2-SAE-Cross** against the ESM2 Base model across the five functional categories. Points above the diagonal indicate datasets where the multi-task gating approach yields higher prediction accuracy than the baseline.

The **ESM2-SAE-Cross** framework achieved state-of-the-art (SOTA) results on 17 datasets within the VenusMutHub benchmark, consistently outperforming established top-tier models including SaProt, VenusREM, Tranception, MIF, and PoET (Figure 4b). To systematically quantify the benefits of cross-task learning, we established **ESM2-SAE-Task** as a rigorous control group. Unlike the cross-fitness model, **ESM2-SAE-Task** is trained and evaluated exclusively on datasets belonging to a single fitness category. This setup intentionally isolates the model from any potential knowledge transfer across different functional domains, establishing a baseline for isolated task supervision.

A systematic comparison between the base ESM-2, the **ESM2-SAE-Task** control group, and **ESM2-SAE-Cross** powerfully reveals the unique advantages of data diversity (Figure 4c). Most notably, in the *small_molecule_binding* category, **ESM2-SAE-Cross** dramatically improved the average Spearman *ρ* from the base ESM-2’s 0.1869 to 0.5028, representing a remarkable increase of 169.0%. Intriguingly, the single-task **ESM2-SAE-Task** model exhibited a significant performance decline in this same category, likely due to the limited sample size and the inherent complexity of the small-molecule binding manifold causing severe overfitting. This stark contrast underscores that training on a diverse pool of fitness data—even if sourced from entirely different functional domains—can enhance the model’s ability to localize fundamental biophysical features that are otherwise unattainable through single-task supervision. This synergistic effect highlights a highly promising direction for supervised variant effect prediction: leveraging cross-task knowledge to capture more robust and generalizable biological signals.

Despite these advancements, **ESM2-SAE-Cross** currently encounters notable challenges in specific datasets where performance remains suboptimal, occasionally falling below the zero-shot ESM-2 baseline (Figure 4d). These discrepancies may stem from variations in experimental conditions, noise in fitness quantification methods, or inherent domain shifts between different deep mutational scanning (DMS) assays. The attempt to globally localize features across highly diverse datasets can potentially lead to cross-dataset overfitting, where certain features identified as universally critical in the training set fail to generalize or even introduce interference in the test set. For instance, a feature associated with binding affinity in one context may act as a confounding factor for protein stability in another. These observations underscore the persistent complexity of the fitness landscape and suggest that while multi-task modeling offers a path toward universal protein understanding, the precise alignment of experimental noise and domain-specific feature importance remains a critical hurdle for future development.

## 3 Discussion

In this study, we introduced a mechanistic intervention framework based on Sparse Autoencoders (SAEs) to address the representation entanglement in protein language models (PLMs). Our results demonstrate that disentangling the latent manifold into discrete, interpretable features significantly enhances variant effect prediction (VEP) across diverse biological contexts.

### Advantages and Generalizability

A primary advantage of the proposed framework is its computational efficiency and architectural simplicity. The training process for the ReLU-SAE is straightforward, utilizing standard reconstruction and sparsity objectives. During the inference phase, the mechanistic intervention—consisting of an encoder mapping, element-wise boosting or gating, and a decoder reconstruction— adds negligible computational overhead. This allows the framework to enhance the zero-shot capabilities of the underlying PLM or be adapted into a supervised gating system without retraining the massive backbone model. Furthermore, our experiments confirm that this approach is generalizable across different PLM architectures, including both the ESM-2 and ESM-3 series.

### Potential for Broader Functional Prediction

The utility of the SAE-based intervention extends beyond variant effect prediction. Because our framework operates directly on the final hidden layers of the PLM, it has the potential to be applied to any protein function prediction task that utilizes PLM-derived embeddings. As demonstrated through the VenusMutHub benchmarks, the framework effectively handles diverse fitness landscapes, including binding affinity, enzymatic activity, and protein stability. This suggests that the disentangled latent space can serve as a universal foundation for various downstream biological tasks.

### Limitations and Future Work

Despite its performance gains, the current framework possesses certain limitations that warrant further investigation. First, the importance of specific SAE features can vary significantly across different protein families or structural environments. Relying on a global localization strategy for critical features may lead to suboptimal results or interference on specific datasets where the “universal” features are less relevant. Second, while we observe substantial performance improvements on specific targets, such as the human E3 ubiquitin ligase HECD1, our current exploration of the underlying biological causes remains limited. We have yet to conduct exhaustive case studies to determine the precise biophysical reasons why the method succeeds or fails on individual proteins. Investigating these failure modes and identifying protein-specific feature importance is a critical direction for future development.

### Bridging Interpretability and Prediction

While pioneering studies have already begun to map the sparse latent space of PLMs to biologically interpretable features [15–17], a significant gap remains in utilizing these insights to actively guide predictions. The future of this field lies not merely in identifying “what” a feature represents, but in understanding “how” these identified mechanisms—such as specific steric clashes or hydrophobic disruptions—can explain the model’s VEP outputs. Beyond explanation, the ultimate goal is to leverage this biological interpretability to further enhance predictive power. By integrating prior human knowledge about specific biochemical constraints into the intervention process, we can move towards a more robust and “knowledge-augmented” paradigm for predicting protein fitness in evolutionary “dark zones”.

## 4 Methods

### 4.1 Architecture and Training of Sparse Autoencoders

#### Data Acquisition and Pre-processing

The foundation of our mechanistic intervention framework rests on a large-scale corpus of protein representations curated from the Protein Data Bank (PDB, accessed as of January 24, 2025) [20] and the Swiss-Prot subset of the AlphaFold Database (AlphaFoldDB) [21]. To facilitate the high-throughput training of Sparse Autoencoders (SAEs), we did not store raw sequences or structural metadata in the training database. Instead, we utilized the high-dimensional embeddings extracted via the base protein language models, which were stored in a Lightning Memory-Mapped Database (LMDB). This architecture ensures constant-time indexing and minimizes I/O overhead during the SAE optimization process by providing direct access to the pre-computed latent manifold.

#### Protein Language Model and Representation Extraction

We utilized two state-of-the-art protein language models (PLMs) as the base encoders for feature extraction: ESM-3 (specifically the 1.4B open variant) and the 650M-parameter variant of ESM-2. For both models, dense embeddings were strictly extracted from their respective final hidden layers (layer 48 for ESM-3 1.4B, and layer 33 for ESM-2 650M). The primary rationale for targeting the final hidden layer is that it directly controls the generation of the log-likelihood ratio (LLR) output. By operating on the representations immediately preceding the final likelihood projection, the extracted embeddings are optimally suited for isolating and disentangling the key mechanistic features that are directly associated with variant effect prediction. For ESM-2, the extraction process was optimized for high-performance computing clusters using a dynamic batching strategy, where the number of sequences per batch was constrained by a maximum token limit (`MAX_TOKENS_PER_BATCH` = 12,000) to maximize NPU/GPU utilization without memory overflow. For ESM-3, we employed a parallel inference pipeline across multiple Huawei Ascend NPU devices to process over two million protein chains, extracting the continuous hidden manifold which served as the input *x* ∈ ℝ^*d*^ for the subsequent SAE models.

#### Sparse Autoencoder Architecture

To disentangle the dense, entangled PLM embeddings into interpretable biological features, we implemented a ReLU-based Sparse Autoencoder (ReLU-SAE). Given an input embedding *x* ∈ ℝ^*d*^, the encoder maps it to a high-dimensional sparse activation space *z* ∈ ℝ^*F*^, where *F* ≫ *d*. The encoder transformation is defined as:

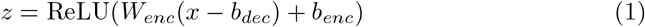

where *W*_*enc*_ ∈ ℝ^*F ×d*^ is the encoder weight matrix, *b*_*enc*_ ∈ ℝ^*F*^ is the encoder bias, and *b*_*dec*_ ∈ ℝ^*d*^ is the pre-encoder bias used to center the data. The hidden representation *z* is then reconstructed back into the original embedding space via a linear decoder:

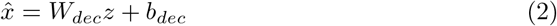

where *W*_*dec*_ ∈ ℝ^*d×F*^ represents the decoder dictionary. For ESM-3 and ESM-2 (650M), we set the activation dimension *d* to 1536 and 1280, respectively, and utilized an expansion factor of 32, resulting in a dictionary size *F* of approximately 40,960 features.

#### Training Objectives and Constrained Optimization

The SAE was trained to minimize a composite loss function ℒ consisting of a reconstruction error and a sparsity-inducing *L*_1_ penalty:

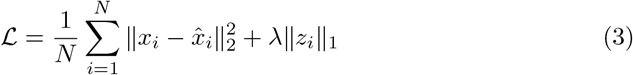

where *N* is the number of residue embeddings in a batch and *λ* is the sparsity coefficient (set to 0.04). To prevent the “feature shrinking” problem and ensure a stable latent manifold, we employed a specialized optimizer, `ConstrainedAdam`. This optimizer maintains unit-norm constraints on the decoder weights *W*_*dec*_ such that each feature vector (dictionary atom) *w*_*j*_ satisfies ∥*w*_*j*_∥_2_ = 1. During each optimization step, `ConstrainedAdam` projects the gradients away from the parallel component of the decoder directions and subsequently renormalizes the weights:

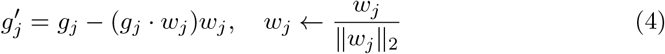

The models were trained for 100,000 steps using a learning rate of 1 × 10^−3^ and a linear warmup period of 500 steps. We utilized a batch size of 40 protein chains, which corresponds to approximately 6,000 residue embeddings per optimization step. This configuration balances the stability of the gradient estimates with the computational efficiency required for large-scale dictionary learning on NPU hardware.

### 4.2 Unsupervised Feature Localization and Zero-Shot Variant Effect Prediction

#### Heuristic Filtering via Marginal Likelihood

The first phase of the **ESM3-SAE-Zero** framework involves identifying a precise subset of high-confidence pathogenic mutations to serve as structural probes. To avoid relying on external labels, we utilize the base ESM-3 model’s zero-shot log-likelihood ratio (LLR) as an unsupervised heuristic filter. For a given wild-type sequence *x* and a mutation at position *i* changing the amino acid to 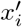, the masked marginal LLR is computed as:

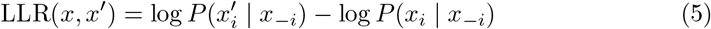

where *x*_−*i*_ represents the masked sequence context. To efficiently navigate the combinatorial explosion of all possible missense mutations across the proteome, we designed a rigorous two-stage filtering pipeline. First, at the individual protein level, we evaluate LLRs for all possible variants but exclusively retain the Top-*K* mutations (empirically set to *K* = 10) exhibiting the lowest LLR scores per chain. We also record the absolute minimum LLR for each respective chain. Second, at the global dataset level, we apply a strict cohort-level filter, retaining only those protein chains whose minimum LLR falls below the 50th percentile across the entire database. This dual-filtering strategy yields a highly enriched probe pool comprising only the most severe mutations from the most mutationally sensitive proteins, essentially isolating the variants that most severely violate the model’s learned structural priors.

#### Mechanistic Feature Localization in the Sparse Manifold

Having isolated the highly-confident pathogenic probes, we sought to mechanically localize the origins of their predicted instability. For each identified probe pair (wild-type *x* and mutant *x*′), we extracted their corresponding final-layer dense embeddings and mapped them into the sparse latent space using the frozen **ESM3-SAE-Zero** encoder. This yielded the high-dimensional sparse activation vectors *z*_*W T*_ and *z*_*MT*_.

To isolate the specific feature dimensions perturbed by the mutation, we employed a contrastive difference metric. The absolute activation difference for each feature dimension *j* was calculated as:

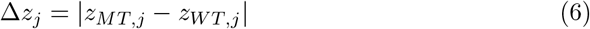

We aggregated these absolute differences across the entire filtered pool of pathogenic probes. By ranking the statistically aggregated Δ*z* scores, we identified the Top-*N* feature dimensions (where *N* = 5000) that exhibited the most acute sensitivity to pathogenic mutations. These high-variance dimensions represent the core mechanistic constraints—such as steric clashes, charge repulsion, or hydrophobic core disruptions—that universally dictate protein fitness.

#### Feature Intervention and Zero-Shot Prediction

With the core pathogenic features mechanically localized, we implemented a direct feature intervention strategy to enhance the zero-shot variant effect prediction (VEP) for novel mutations. During inference, the masked input sequence of a target variant is processed to obtain its mutant sparse representation *z*_*MT*_. We then apply an element-wise boosting mask *M*_*boost*_ ∈ ℝ^*F*^ to selectively amplify the localized Top-*N* pathogenic features. The intervened sparse representation 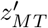 is formalized as:

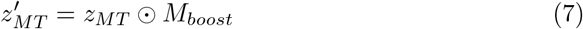

where *M*_*boost,j*_ = *γ* (e.g., boosting factor *γ* = 2.0) if dimension *j* belongs to the identified Top-*N* core features, and 1.0 otherwise.

Finally, the intervened sparse vector is reconstructed back into the dense continuous space via the SAE decoder:

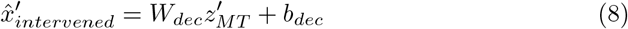

This enhanced dense representation, which now carries an amplified signal for critical biophysical constraints, is subsequently passed back to the frozen ESM-3 Language Model (LM) head. The resulting output probabilities are used to compute the intervened LLR, establishing a structurally-aware and highly sensitive metric for zero-shot variant effect prediction without requiring any task-specific fine-tuning.

#### Algorithm Flow

The complete unsupervised pipeline, encompassing heuristic filtering, feature localization, and the zero-shot mechanistic intervention, is formally outlined in Algorithm 1.

##### Algorithm 1

Unsupervised Feature Localization and Zero-Shot Intervention (**ESM3-SAE-Zero**)

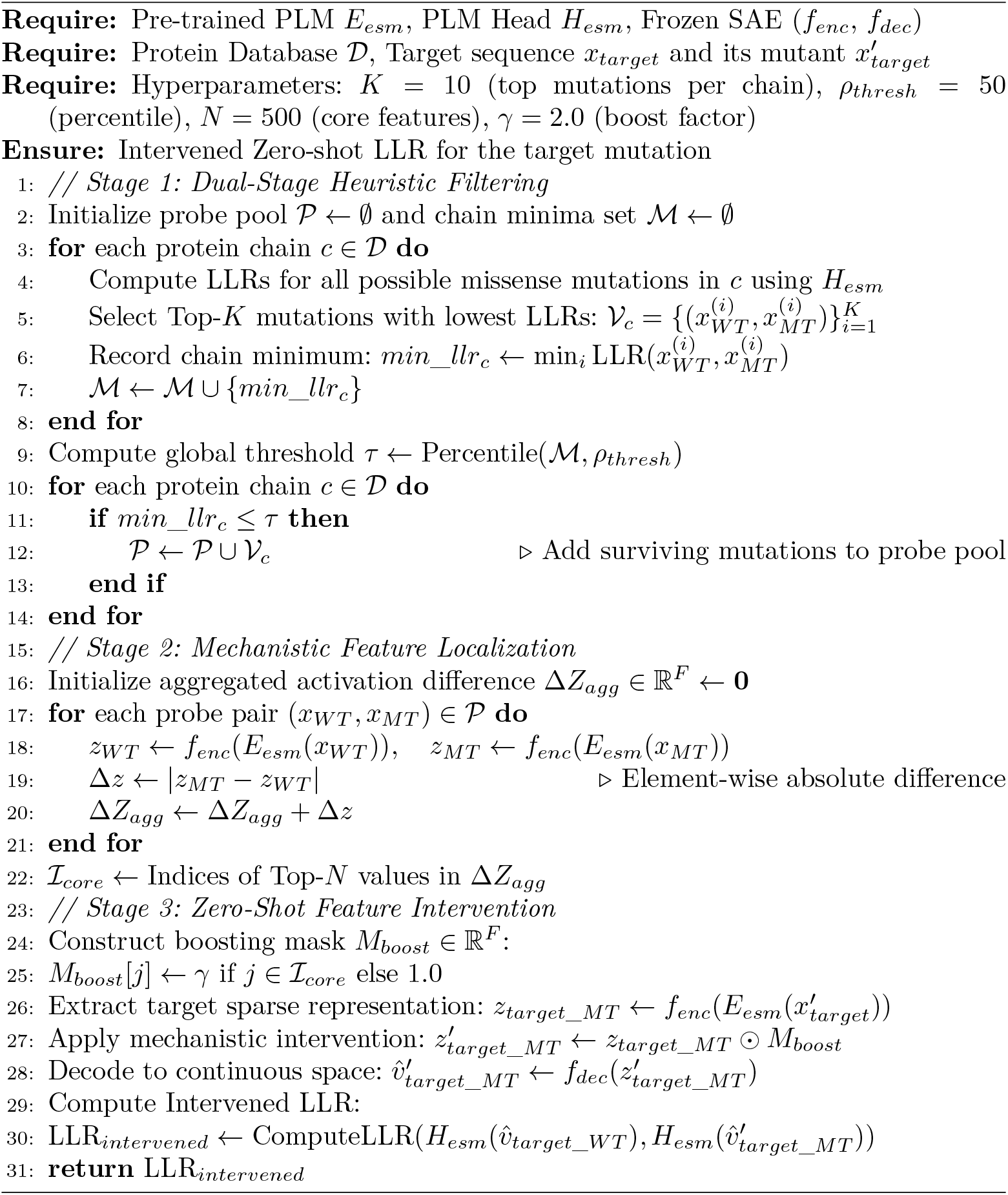

### 4.3 Target-Specific Differentiable Gating and Optimization

#### Differentiable Feature Masking via Straight-Through Estimator

While the unsupervised approach (**ESM3-SAE-Zero**) identifies universally critical features, optimizing predictions for distinct biological targets requires isolating the specific submanifold of the latent space relevant to a particular functional assay. To achieve this, we introduce the **ESM3-SAE-Target** model, which incorporates a trainable binary gating module operating on the frozen sparse latent space of the SAE.

For each protein target, we define a trainable parameter vector *s* ∈ ℝ^*F*^ representing the unnormalized logits of the gate. To perform hard feature selection while maintaining end-to-end differentiability, we utilize a Straight-Through Estimator (STE). During the forward pass, the binary mask *m* ∈ {0, 1}^*F*^ is generated by thresholding the sigmoid probabilities *σ*(*s*) = 1*/*(1 + exp(−*s*)):

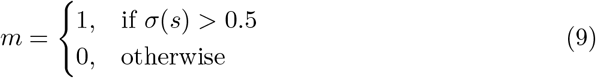

The gated sparse representation used for reconstruction is then defined as *z*_*gate*_ = *z* ⊙ (*m* + *σ*(*s*) − sg[*σ*(*s*)]), where sg[·] denotes the stop-gradient (detach) operator. This formulation ensures that in the forward pass, the model uses a discrete binary mask for feature intervention, while in the backward pass, the gradient of the loss ℒ with respect to the gate logits is approximated via the continuous sigmoid function:

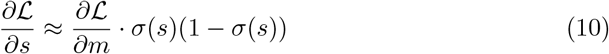

This mechanism allows the framework to learn an optimal, target-specific binary configuration of the latent manifold through gradient-based optimization.

#### Optimization Objective and Sparsity Constraint

The gated sparse vector is reconstructed back into the dense continuous space and subsequently passed to the frozen ESM-3 sequence prediction head to compute the predicted log-likelihood ratio (LLR_*pred*_). To optimize the gate *s* for variant effect prediction, we employ a supervised learning strategy. The primary objective is to maximize the rank correlation between the predicted LLRs and the experimental fitness scores (*y*) from deep mutational scanning (DMS) datasets. We utilize a Soft-rank Pearson correlation loss, ℒ_*rank*_, which employs a differentiable soft-sorting operator to provide a continuous approximation of the Spearman rank correlation.

To ensure the model identifies a parsimonious set of mechanistic features and avoids overfitting to the entangled dense embeddings, we incorporate an *L*_1_ sparsity penalty on the gate probabilities. The total loss function minimized during training is:

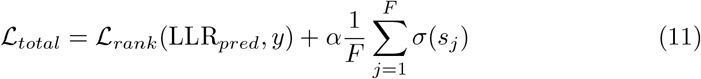

where *α* is a regularization hyperparameter (typically set to 1.0) that balances predictive accuracy with feature sparsity.

#### Cross-Validation and Training Procedure

The **ESM3-SAE-Target** gate is optimized for each individual protein target using the AdamW optimizer. To ensure the robustness and generalizability of the identified features, we follow the official ProteinGym benchmarking protocol, employing a five-fold random cross-validation split. In each fold, the gating module is trained on the training mutations and evaluated on the held-out test set. The final performance reported is the average Spearman *ρ* across all folds. By restricting the supervised signal to the selection of pre-trained sparse features rather than fine-tuning the entire model, the framework maintains the foundational biophysical knowledge of the PLM while successfully adapting to the unique functional landscape of the specific target.

### 4.4 Multi-Task Cross-Fitness Architecture and Regularization

#### Modular Multi-Task Gating Mechanism

To enable simultaneous training across highly heterogeneous fitness landscapes (e.g., stability, binding, activity) within the VenusMutHub benchmark, we designed the **ESM2-SAE-Cross** architecture. This unified framework utilizes a decoupled shared-private gating mechanism to dynamically route sparse features. For a sparse latent space of dimension *F* and a set of *T* distinct biological tasks, we define a single shared gate parameterized by unnormalized logits *s*_*shared*_ ∈ ℝ^*F*^, and a set of task-specific private gates parameterized by 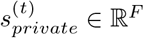 for each task *t* ∈ {1, …, *T*}.

The shared gate is designed to capture universal biophysical constraints (e.g., backbone steric clashes or fundamental hydrophobic packing), while the private gates capture domain-specific functional requirements. Similar to the single-target model, we generate continuous probabilities via the sigmoid function: *p*_*shared*_ = *σ*(*s*_*shared*_) and 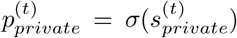. To maintain end-to-end differentiability while enforcing discrete feature selection, we apply the Straight-Through Estimator (STE) to obtain the binary masks *b*_*shared*_ and 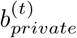.

For an input sequence belonging to task *t*, the representations from the shared and private manifolds must be combined. We accomplish this through a differentiable logical OR operation, mathematically formulated using a clamped addition:

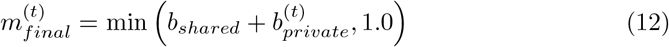

The mutant sparse representation is then selectively activated via 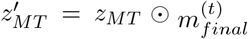. This parallel structure ensures that a sparse feature is preserved if it is deemed critical by either the universal shared constraint or the task-specific requirement, preventing catastrophic forgetting across diverse datasets.

#### Objective Functions and Orthogonal Regularization

A critical challenge in multi-task feature routing is preventing the collapse of private gates into the shared gate, ensuring that the network distinctly allocates universal features to the shared manifold and specialized features to the private manifolds. The total optimization objective for a mini-batch of task *t* is composed of the empirical rank loss (ℒ_*rank*_) and a set of structural regularizers:

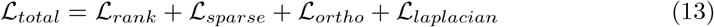

First, we utilize the Soft-rank Pearson correlation loss (ℒ_*rank*_) to align the predicted LLRs with the true experimental fitness values. Second, to enforce a parsimonious feature selection across the entire architecture, we apply a global *L*_1_ sparsity penalty to both the shared and private continuous probabilities:

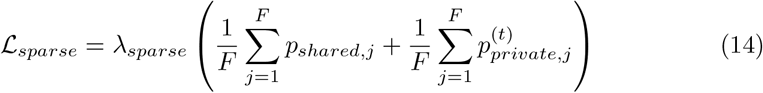

Third, to forcefully disentangle the universal and task-specific features, we introduce an orthogonality loss (ℒ_*ortho*_). This term penalizes the intersection (dot product) between the shared and private probabilities, explicitly discouraging them from concurrently activating the same sparse feature dimension:

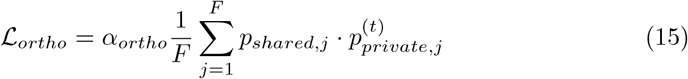

#### Feature Co-activation Graph Laplacian

A critical innovation in our framework is the incorporation of a Graph Laplacian regularization (ℒ_*laplacian*_) to capture the inherent collaborative relationships between biological features. During pre-training, we empirically observed that certain SAE features frequently co-activate across diverse protein families, implying that they act synergistically to form stable structural motifs. To encode this structural prior, we pre-computed a dense feature co-activation matrix *A*_*dense*_ ∈ ℝ^*F ×F*^ by aggregating the global maximum pooling of binary feature activations across the entire pre-training corpus.

We define a hard-thresholded adjacency matrix *A* where *A*_*i,j*_ = *A*_*dense,i,j*_ if *A*_*dense,i,j*_ *> τ* (e.g., *τ* = 0.05), and 0 otherwise. Let *D* be the diagonal degree matrix where *D*_*i,i*_ = ∑_*j*_ *A*_*i,j*_. The Graph Laplacian is defined as *L*_*graph*_ = *D* − *A*. To encourage the gating mechanism to respect this topological prior—such that synergistically co-activating features are selected or pruned jointly—we apply the Laplacian penalty to both the shared and private gate probabilities via their quadratic forms:

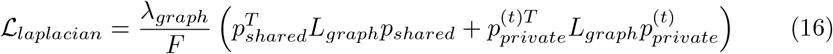

Computationally, this is efficiently resolved without explicitly instantiating *L*_*graph*_ by calculating the degree and edge terms directly: *p*^*T*^ *L*_*graph*_*p* = *p*^*T*^ (*D* ⊙ *p*) − *p*^*T*^ (*Ap*). This regularization dynamically smooths the decision boundaries in the latent space, forcing the model to select functionally cohesive sub-networks rather than isolated features.

#### Cross-Fitness Inference

During forward inference, the mutated sequence is processed by the frozen ESM-2 (650M) backbone. The combined gated sparse features 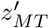 are reconstructed back into the dense 1280-dimensional manifold via the SAE decoder and fed directly into the ESM-2 Language Model (LM) head. The multitask framework is trained jointly by shuffling mixed batches from all VenusMutHub categories. This unified cross-fitness approach, constrained by explicit biophysical co-activation priors, circumvents the data sparsity limitations of individual assays, yielding highly robust and generalizable variant effect predictions.

### 4.5 Downstream Evaluation Benchmarks and Protocols

#### ProteinGym Benchmark and Cross-Validation Protocol

To rigorously evaluate the zero-shot mechanistic intervention (**ESM3-SAE-Zero**) and the target-specific gating framework (**ESM3-SAE-Target**), we utilized the ProteinGym benchmark. ProteinGym is an expansive and highly standardized compendium of deep mutational scanning (DMS) assays, encompassing a wide array of protein families, taxonomic groups, and functional mechanisms. Our evaluation strictly included the 217 substitution datasets that provide high-quality experimental fitness measurements.

For the supervised **ESM3-SAE-Target** model, preventing data leakage and overfitting is paramount. Consequently, we adhered strictly to the official ProteinGym 5-fold random split protocol (`fold_random_5`). For each dataset, the differentiable feature gate was optimized exclusively on the training folds (80% of the mutations) utilizing the Soft-rank Pearson correlation loss, and subsequently evaluated on the held-out testing fold (20% of the mutations). This process was repeated across all five folds, and the final reported performance for a given protein target is the average of the evaluation metrics across all folds. Spearman’s rank correlation coefficient (*ρ*) was utilized as the primary metric to assess the monotonic relationship between the predicted log-likelihood ratios (LLRs) and the experimental fitness scores.

#### VenusMutHub for Cross-Fitness Evaluation

Evaluating the generalization and knowledge-transfer capabilities of our multi-task architecture (**ESM2-SAE-Cross**) requires a benchmark that explicitly disentangles distinct biological mechanisms. Therefore, we employed the VenusMutHub benchmark, which categorizes DMS datasets into a granular taxonomy of five distinct fitness classes: *activity, ppi_binding, selectivity, small_molecule_binding*, and *stability*.

This explicit categorization is critical for our cross-fitness experimental design for two reasons. First, it allows us to establish rigorous isolated single-task controls (**ESM2-SAE-Task**), where a model is strictly trained and evaluated within a single functional domain (e.g., only on *stability* datasets). Second, it provides the structural foundation for our decoupled shared-private gating mechanism, allowing the network to route universal biophysical constraints into the shared gate while allocating domain-specific features to the categorical private gates. During the evaluation of the multi-task framework, datasets within each of the five functional categories were split into training and evaluation sets. The model was optimized jointly across the heterogeneous training sets and evaluated on the unseen test sets, enabling us to quantify the synergistic performance gains achieved through cross-fitness feature co-activation.

#### Baseline Comparisons and SOTA Evaluation

For a rigorous and standardized assessment against existing state-of-the-art (SOTA) models, we utilized the official evaluation scores explicitly provided by the ProteinGym and VenusMutHub repositories. The performance metrics (e.g., Spearman *ρ*) for all baseline architectures—including but not limited to SaProt, VenusREM, Tranception, MIF-ST, and PoET—were directly sourced from the official dataset releases and leaderboard data, rather than through independent reimplementation. This protocol ensures that all competitor models are represented by their optimally tuned, officially reported capabilities, eliminating potential discrepancies caused by sub-optimal hyperparameter tuning during third-party reproduction. Consequently, the calculations for average performance gains (Δ*ρ*) and head-to-head win rates presented in our results are strictly grounded in these verified, publicly accessible benchmark scores.

## Notes

### Competing Interest Statement

The authors have declared no competing interest.

